# A universal set of primers to study animal associated microeukaryotic communities

**DOI:** 10.1101/485532

**Authors:** Javier del Campo, Maria J. Pons, Maria Herranz, Kevin C. Wakeman, Juana del Valle, Mark J. A. Vermeij, Patrick J. Keeling

## Abstract

**Background:** Unlike the study of bacterial microbiomes, the study of the microeukaryotes associated with animals has largely been restricted to visual identification or molecular targeting of particular groups. The application of high-throughput sequencing (HTS) approaches, such as those used to look at bacteria, has been restricted because the barcoding gene traditionally used to study microeukaryotic ecology and distribution in the environment, the Small Subunit of the Ribosomal RNA gene (18S rRNA), is also present in the animal host. As a result, when host-associated microbial eukaryotes are analyzed by HTS, the obtained reads tend to be dominated by host sequences.

**Results:** We have done an in-silico validation against the SILVA 18S rRNA reference database of contrametazoan primers that cover the V4 region of the 18S rRNA, and compared these with universal V4 18S rRNA primers that are widely used by the microbial ecology community. We observe that the contrametazoan primers recover only 2.6% of all the metazoan sequences present in SILVA, while the universal primers recover up to 20%. Among metazoans, the contrametazoan primers are predicted to amplify 74% of Porifera sequences and 4% and 15% of ctenophore and Cnidaria, respectively, while amplifying almost no sequences within Bilateria.

We tested these predictions in-vivo, and observed that contrametazoan primers amplify the 18SrRNA from two ctenophore species, but reduce significantly the metazoan signal from material derived from coral and from human samples. When compared in-vivo against universal primers, contrametazoan primers worked in 8 out of 9 samples, providing at worst a 2-fold decrease in the number of metazoan reads, and at best a 2800-fold decrease.

**Conclusions:** We have validated an easy, inexpensive, and near-universal method for the study of microeukaryotes associated with animal hosts using 18S rRNA Illumina metabarcoding. This method will contribute to a better understanding of microbial communities, as they related to the wellbeing of animals and humans.

## Background

Since the publication of the human microbiome consortia paper [1] there has been an acceleration in the study of host-associated microbes using metabarcoding methods in many different animal systems. While the term “microbe” spans a wide range of organisms, most published studies examining microbiomes focus exclusively on bacteria [2], some focus on fungi [3], and only a few focus on other microeukaryotes (protists) [4]. However, protists are part of a healthy microbiota and play a relevant role in the mammal gut ecosystem (including humans) [5], altering the diversity and composition of the gut communities as a whole [6], interacting directly with the host immune system, and even contributing to mucosal immunity conferring disease protection [7]. In other animals, beyond mammals, microeukaryotes and protists in particular contribute to key host processes, like cellulose digestion [8], and in some cases, like the zooxanthellae in corals, microeukaryotes are crucial for the survival of the host [9]. Apart from these many beneficial roles, microeukaryotes are also well-known parasites, and their action can have dramatic effects on the fitness of the host [10]. So why have they been largely ignored from microbiome studies for so long?

There are several reasons for this. For one, bacteria are, in general, the primary or only focus of most microbiological work, not only in host-associated environments but also in other systems [11]. But in the case of host-associated microeukaryotes there is an additional technical issue that complicates the study of microeukaryotes using metabarcoding. The most common markers used for metabarcoding are 16S rRNA for bacteria, ITS for Fungi, and 18S rRNA for the eukaryotes. All of them have been successfully applied in free-living environments but their success differs when used in animal host-associated environments. The main issue is that when using universal primers, most of the sequences recovered from PCR using universal 18S primers are from the host itself and not its microeukaryotic microbiome [12–14], sometimes by many orders of magnitude. One solution to this problem is to use blocking primers in the PCR reaction, which are chemically modified primers (with a C3 spacer) that target the host 18S rRNA and will prevent the extension during the PCR when using universal primers [15]. But because metazoans are so diverse, blocking primers have to be specifically designed case by case for each host [16–20], so this might work for a subject of intensive study like humans, but not for any broader study of animal microbiome diversity. Another option is to use specific primers for eukaryotes that avoid a particular host [21], this approach seems to be easier than the use of blocking primers, but once again requires a specific set of primers for every metazoan group. Similarly, one particular lineage of eukaryotes can be investigated using primers specific for that group and excluding animals (like those used for fungi), but in this case, no other aspect of the eukaryotic diversity can be assessed.

Our aim was to test a more universally applicable approach to recover the eukaryotic component of the microbiome, the eukaryome [22] from as many animals as possible, and in the easiest and least expensive way possible. To this end, we screened the parasitology literature for primers that have been used to detect parasites in animals, and we focused on studies that screened for parasites in a wide range of hosts. In one study, by Bower and colleagues in 2004 [23], they assessed primers in “mock communities” consisting of metazoan tissue mixed with parasites from culture, and identified a pair of primers that they defined as universal non-metazoan primers (UNonMet). These primers allowed Bower et al. to screen for several kinds of parasites in a broad range of metazoans, from mollusks, to nematodes, to fish. We considered these primers a potential candidate for the study of the eukaryome as a whole using a metabarcoding approach and here test them in-silico and in-vivo.

## Results

Based on the results of Bower et al. [23] and our own amplifications in the lab, the product of UNonMet primers was typically 600bp, and when aligned against the *Saccharomyces cerevisiae* 18S rRNA gene, the amplicon covers the V4 and flaking regions. We first tested the UNonMet primer in-silico against the SILVA 18S rRNA Reference Database [24]. The SILVA database contains all the 18S rRNA sequences longer than 1200 bp present in GenBank and it is considered a reliable source for probe and primer design. Testing the UNonMet primers against SILVA recovered only 2.6% of the metazoan 18S reads that were present, whereas they still performed well with the rest of the Eukaryotic groups, except among excavates, which will be discussed below (Fig 1). Examining the distribution of the metazoan diversity that was recovered in more detail, reveals that most of the reads came from sponges - the primers recover up to 74% of the sponge reads present in Silva - as well as a small number of ctenophores (4%) and cnidarians (15%). Fewer than 1% of reads from bilaterians are recovered (Fig 2).

**Figure 1.**
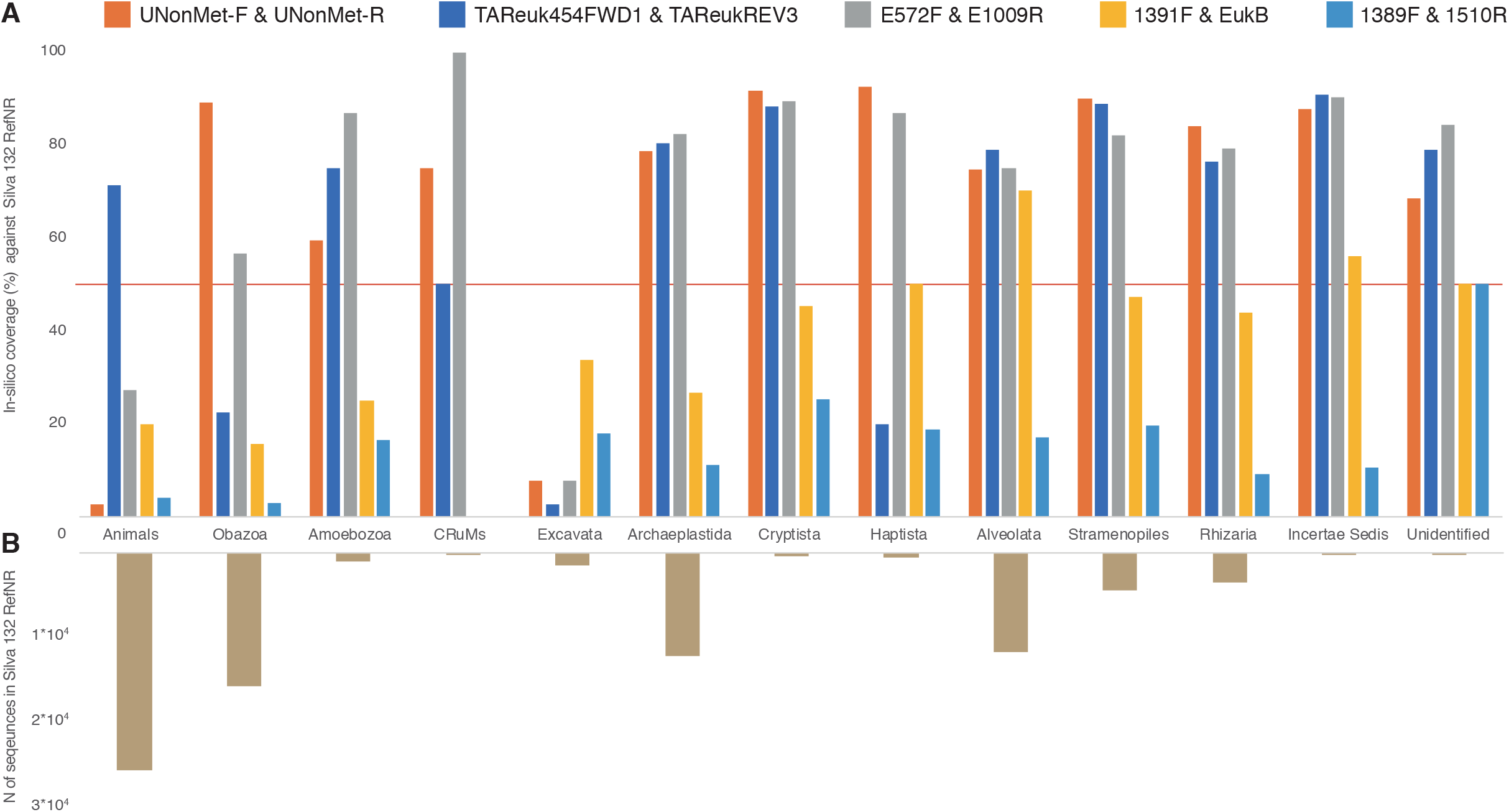
Comparison the in-silico 18S rRNA diversity recovered from SILVA 132 RefNR using UNonMet primers versus the most used V4 and V9 18S rRNA primers for microeukaryotic metabarcoding. A) Percentage of recovered diversity from the metazoans and each of the major eukaryotic groups B) Number of reads for each group present in SILVA 132 RefNR

**Figure 2.**
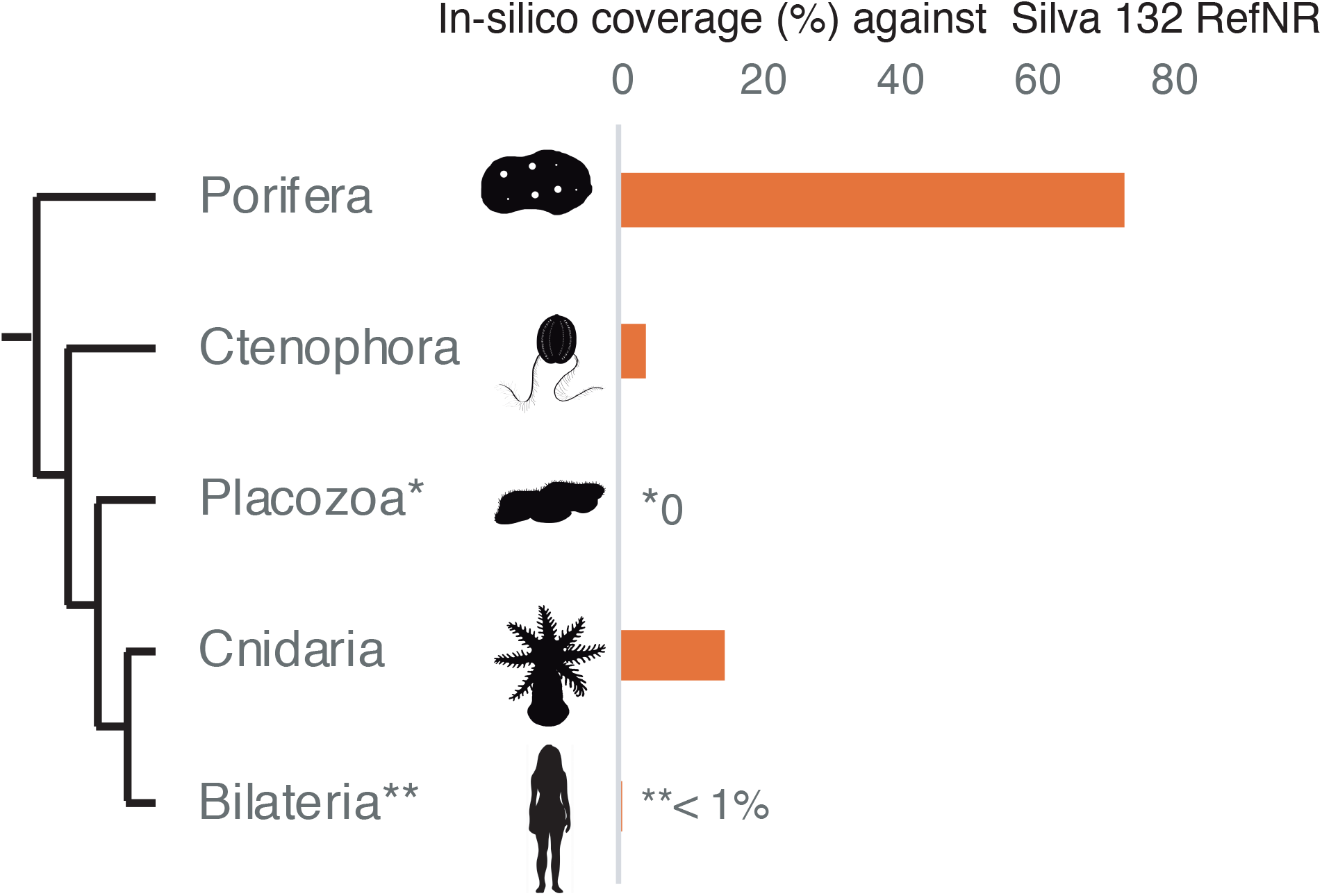
Percentage of in-silico metazoan18S rRNA diversity recovered from SILVA 132 RefNR using UNonMet primers. A more detailed analysis at Phylum level is available at Supplementary Table 1.

Next, we compared these results with in-silico tests of the most commonly used primers in microeukaryotic metabarcoding (Table 1). For the comparison we selected 4 sets of universal 18S primers, based on their relevance in current work and potential to limit metazoans. TAReuk454FWD1 and TAReukREV3 amplify the V4 region and were used in the European coastal study BioMarKs [25,26] and the global ocean survey Malaspina [27], and have subsequently become the most widely used V4 primers; E572F and E10009R [28] are the V9 universal primers suggested by the Earth Microbiome Project [29]; 1391F and EukB also amplify the V9 region and were used by the Tara Oceans Consortium [30]. Both sets of V9 primers are widely used. Additionally, we compared a set of V4 universal primers E572F and E10009R developed by Comeau et al. [31] as V4 universal primers that recover fewer metazoan reads. Focusing on the percentage of metazoans reads recovered, the UNonMeta primers recover the fewest and TAReuk recovers the most. The Tara Oceans primers also recover relatively few metazoans, however, their performance with the rest of eukaryotic groups is poor, failing to recover more than 30% of the diversity within most of groups of protists (Fig 1). Within the rest of the eukaryotes the UNonMet primers perform as well or in some cases even better that the analyzed universal primers, except in the case of the excavates, where its performance is poor. This is generally the case with V4 universal primers, and has been previously reported as a known limitation [32].

**Table 1.**
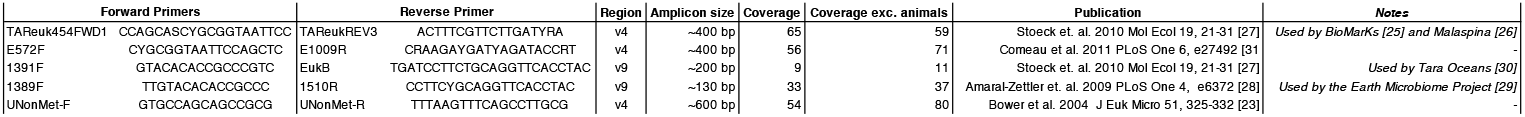
List of primers tested, region, amplicon size, publication and SILVA 132 RefNr coverage.

To see whether these in-silico analyses reflected real performance in-vivo, we tested the UNonMet primers on a range of animal host-associated communities. The expected amplicon size (600bp) is at the high end of the size limit that the latest MiSeq Illumina technology can handle in the best-case scenario (2 x 300 bp). Usually the quality of the sequences at the end of the amplicon pairs tends to be low and those fragments are excluded, so the realistic amplicon size of MiSeq reads in typically around 500 bp (2 x 250bp). To enable the merging of paired-ends, we therefore used a two-step approach to proceed with the in-vivo test. We amplified the samples with the UNonMet primers in order to reduce the metazoan signal present, and reamplified these products using the primers described by Comeau et al. [31] that only retrieve 20% of metazoans. The samples analyzed came from Ctenophora, Cnidaria (corals), and Bilateria (humans). We did not examine Porifera because the in-silico results clearly suggest they are unlikely to work. The resulting read distribution shows that the primers did not perform well with the two species of ctenophores analyzed (5 samples), where most of the recovered signal correspondent to the host (Fig 3). In contrast, read mapping from corals (11 samples) and humans (19 samples) resulted in less than 10% of the signal coming from animals (Fig 3), allowing us to detect or increase the signal of various groups of microeukaryotes associated with these metazoan hosts, or other eukaryotic signal likely associated with the diet of the analyzed individuals (most obviously in the high proportion of plant sequences derived from the human samples).

**Figure 3.**
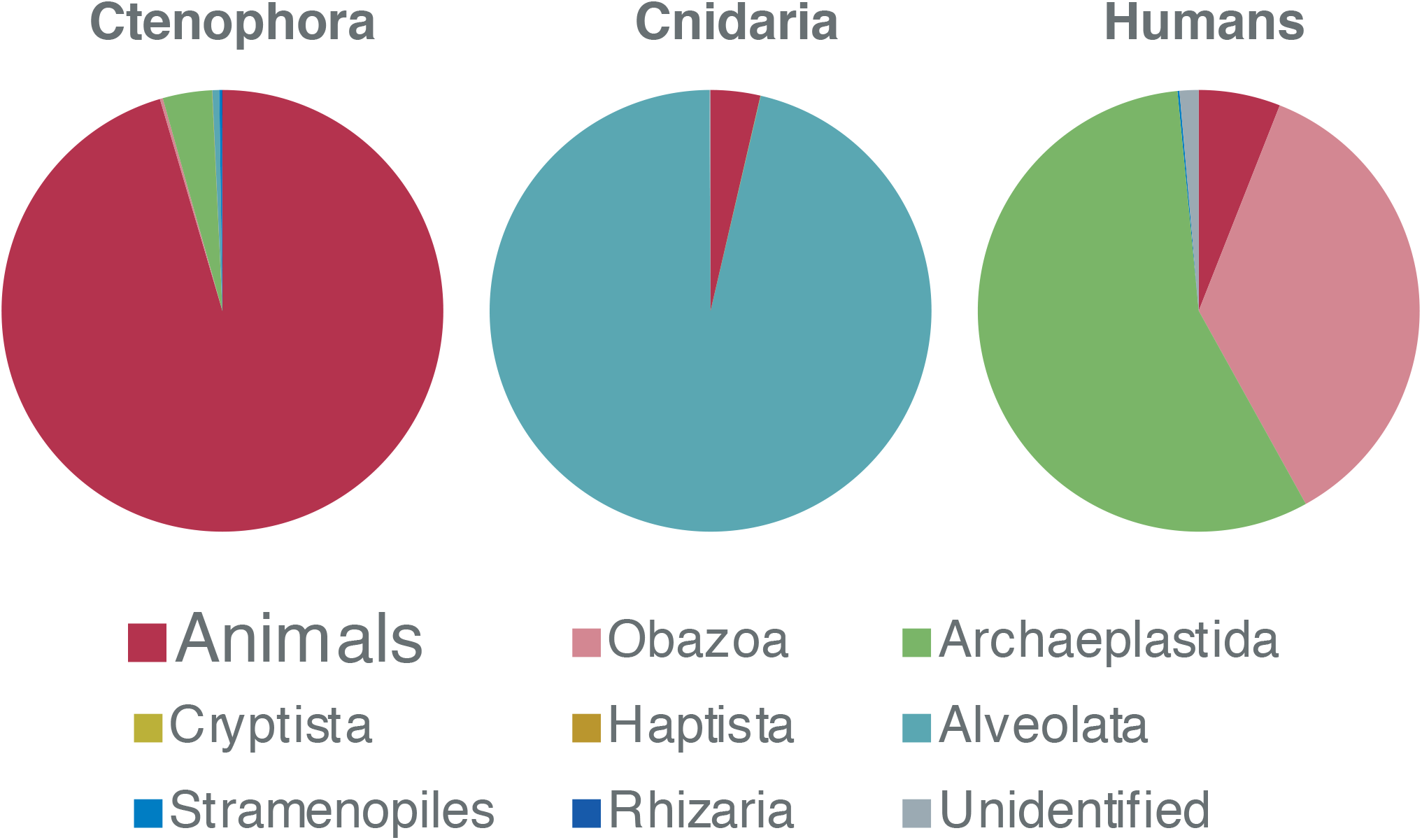
Relative abundance of 18S rRNA reads in Ctenophora, Cnidaria (corals) and Bilateria (Humans) using UNonMet primers.

A subset of the coral and human samples tested using the UNonMet primers were reanalyzed using only the Comeau primers to determine the impact of the contrametazoan primers (Fig 4). In 8 out of 9 compared samples there was a significant decrease in the load of metazoan reads, even in those samples were the initial animal signal was very low. In the worst-case scenario there was a 2-fold reduction of the metazoan reads, whereas the most dramatic reduction was 2,200 times lower.

**Figure 4.**
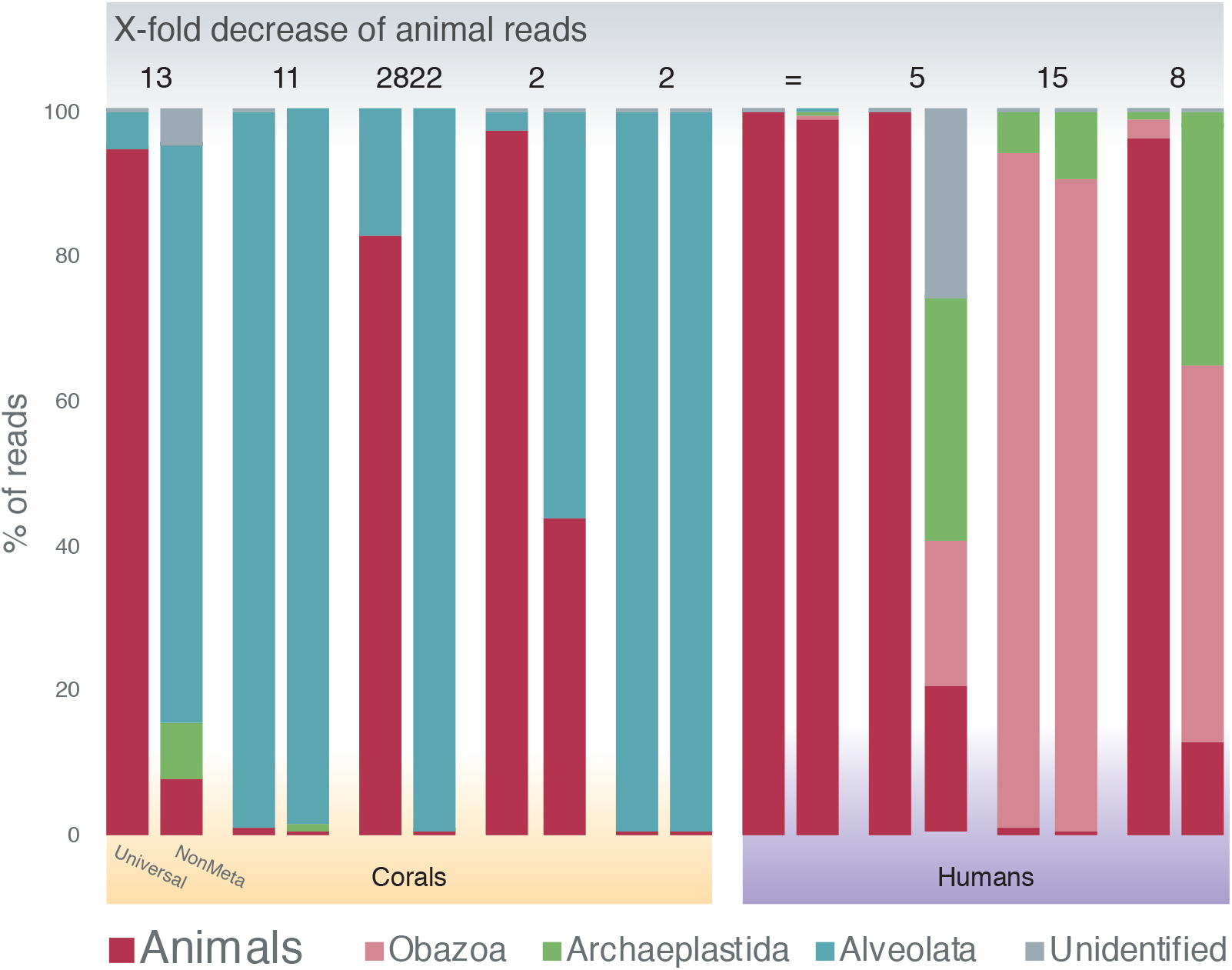
Comparison of the relative abundance of 18S rRNA reads in corals and humans using UNonMet vs Comeu Universal V4 primers.

## Discussion

Both the in-silico analysis and in-vivo results show that the UNonMet primers work well to significantly reduce the metazoan signal from HTS datasets and constitute a suitable approach to access and study the microeukaryotic communities associated with animal hosts. This pair of primers not only excludes the metazoan signal, but they perform as well or sometimes even better than the most commonly used universal 18S rRNA primers in terms of their ability to amplify V4 from a known range of microeukaryotes. Despite the reduction of metazoan signal, there always remains some host signal that we attribute to non-specific amplification because of the amount of host biomass. The degree of this signal is not sufficient to limit the analysis of microeukaryotes, and indeed we see this as an advantage because it gives an independent identification of the animal host, or could indicate if two animals were present in such a case.

The use of contrametazoan primers is clearly an advantage compared with other methods for several reasons. First, it eliminates the need to discard most of the sequence data as is the case universal primers are used alone. It is true that sequencing to greater depth can still allow access to eukaryome diversity, but this is inefficient and expensive. Second, blocking primers have been successfully used to access the eukaryome, but there is no universal blocking primer that will eliminate all or even most animals, so blocking primers tend to be specific to a particular study. The UNonMet primers can be used with most animals, except sponges and perhaps ctenophores, and we show in vivo are effective with corals (cnidarian) and humans (bilaterians), two groups that are far apart in the animal tree of life. Third, the 18S rRNA fragment recovered by UNonMet covers the V4 region, which provides reasonably good phylogenetic resolution, so the barcoding can be more precise than V9-based data. A better phylogenetic resolution makes the results useful not only for microbial ecology, but eventually also for diagnosis. Fourth, this method is both relatively simple and inexpensive, making it broadly available, and the data accordingly better comparable between different studies.

The main concern about this approach is the need for a two-step amplification. This is a limitation because it may lead to additional sequencing errors (although we should note the second round involves limited rounds of amplification so it is not clear if this is a significant problem), and two stages of potential amplification bias. The bias is an issue with all methods, and generally samples analyzed in the same way give a relative picture while samples treated in different ways are hard to compare, and this method is much the same in that regard. However, this is only a limitation because the size of the amplicon generated by UNonMet is slightly larger than can be handled by current Illumina technologies. It is not unreasonable to assume that these limitations will soon be eliminated by changes in the sequencing platforms and chemistry that will allow the UNonMet to be used directly for Illumina libraries (or other chemistries), which would also make the process even simpler and less expensive.

## Conclusions

We have identified and tested a relatively simple solution to an issue that presents a very obvious limitation on microbiome research since the rise of high-throughput metabarcoding was applied to the study of the animal-associated microbial communities. Using the UNonMet primers will allow research on metazoan host-associated environments to more easily include analyses of eukaryome, providing a much fuller picture of the community and finally begin to fill the knowledge gap in microbiome studies of not only humans, but most animals.

## Methods

The in-silico test was done using the TestPrime 1.0 [33] against SILVA 132 RefNR [24] allowing no mismatches.

Corals were collected from several locations in Curaçao between April 19 and 21 2015 and Okinawa (Japan) between April 24 and 26 2015. Whole samples, including skeleton and tissue, were homogenized using mortar and pestle. DNA was extracted with the RNA Powel Soil Total RNA Isolation Kit coupled with the DNA Elution Accessory Kit (MoBio) in the case of the Curaçao samples and DNeasy Blood & Tissue Kit (Qiagen) in the case of the Japan samples. Ctenophores were collected from Calvert Island (British Columbia, Canada) between June 5 and 7 2015. The ctenophore gut and body were extracted and homogenized and DNA was extracted with DNA Power Soil (MoBio). Human stool samples were obtained from children less than 5 years of age who were hospitalized with acute gastroenteritis from January 2011 to April 2014 in the “Hospital Docente Regional de Cajamarca” located in Cajamarca Department, in rural Northern Peru [34]. Genetic material from faecal suspensions were extracted using the High Pure RNA Isolation Kit (Roche Applied Science) in accordance with the manufacturer’s instructions. DNA concentration was quantified on a Qubit 2.0 Fluorometer (Thermo Fisher Scientific Inc.). A complete list of samples is available at Supplementary Table 2.

Eukaryotic microbiome amplicons were prepared using PCR with high-fidelity Phusion polymerase (Thermo Fisher Scientific Inc.), using primers that target the V4 region of 18S rRNA gene, but which exclude metazoan sequences (UNonMetF 5′-GTGCCAGCAGCCGCG-3′, UNonMetR 5′-TTTAAGTTTCAGCCTTGCG-3′) [23]. PCR was performed using the following protocol: 30s at 98°C, followed by 35 cycles each consisting of 10 s at 98 °C, 30 s at 51.1°C and 1 min at 72°C, ending with 5 min at 72°C. PCR products were visually inspected for successful amplification using gel electrophoresis with 1% agarose gels. PCR products were then cleaned using the QIAquick PCR Purification Kit (Qiagen) and quantified on a Qubit 2.0 Fluorometer. Amplicon sequencing was performed by the Integrated Microbiome Resource facility at the Centre for Comparative Genomics and Evolutionary Bioinformatics at Dalhousie University. PCR amplification from template DNA was performed in duplicate using high-fidelity Phusion polymerase. A single round of PCR was done using “fusion primers” (Illumina adaptors + indices + specific regions) targeting the V4 region of the Eukaryotic 18S rRNA gene (primer set E572F + E1009R [31]; ~(~440 bp fragment) with multiplexing. PCR products were verified visually by running a high-throughput Invitrogen 96-well E-gel. The duplicate amplicons from the same samples were pooled in one plate, then cleaned-up and normalized using the high-throughput Invitrogen SequalPrep 96-well Plate Kit. The samples were then pooled to make one library which was then quantified fluorometrically before sequencing on an Illumina MiSeq using a 300 bp paired-end read design.

Amplicon reads were processed (dereplication, chimera detection, and singleton removal) using VSEARCH [35]. Operational taxonomic units (OTU) were clustered at 97% identity using VSEARCH and analyzed using Qiime v1.9.1 [36]. The taxonomic identity of each OTU was assigned based on the SILVA 132 RefNR database [24], using the assign_taxonomy function in Qiime. OTU that were “unassigned” were inspected using BLAST against the GenBank nr database and manually reassigned to the closest hit if possible. OTU represented by fewer than four reads were removed, as were OTU that were identified as metazoan 18S rDNA or mitochondria. Samples with fewer than 1500 reads were excluded from downstream analysis.

## Supporting information

## List of abbreviations

HTS: high-throughput sequencing
18S rRNA: Small Subunit of the Ribosomal RNA gene
UNonMet: Universal non-metazoan primers
OTU: Operational Taxonomic Units

## Acknowledgements

We thank Forest L. Rohwer, Ben Knowles, Emma E. George and Jan Janouškovec for assistance and advice in coral sample retrieval and processing.

## Availability of data and materials

The 18S rRNA amplicon reads are deposited in the European Nucleotide Archive (PRJEB24453, PRJEB29965) and the NCBI Sequence Read Archive (PRJNA482746).

## Authors’ contributions

JdC, and PJK designed the study. JdC, MJP, MH, KW, JdV, MV, and PJK obtained samples. JdC. performed the analyses. JdC wrote the manuscript, with input from all authors. PJK supervised the work.

## Ethics approval and consent to participate

Not applicable

## Consent for publication

Not applicable.

---

**Supplementary Table 1**. Percentage of in-silico metazoan18S rRNA diversity recovered from SILVA 132 RefNR using UNonMet primers at Phylum level.

**Supplementary Table 2**. List of samples including basic taxonomic information and the sampling site.

## References

1. Huttenhower C, Gevers D, Knight R, Abubucker S, Badger JH, Chinwalla AT, et al. Structure, function and diversity of the healthy human microbiome. Nature. Nature Publishing Group; 2012;486:207–14.

2. McFall-Ngai M, Hadfield MG, Bosch TCG, Carey H V., Domazet-Lošo T, Douglas AE, et al. Animals in a bacterial world, a new imperative for the life sciences. Proc Natl Acad Sci. 2013;110:3229–36.

3. Nash AK, Auchtung TA, Wong MC, Smith DP, Gesell JR, Ross MC, et al. The gut mycobiome of the Human Microbiome Project healthy cohort. Microbiome. Microbiome; 2017;5:153.

4. Andersen LO, Vedel Nielsen H, Stensvold CR. Waiting for the human intestinal Eukaryotome. ISME J. 2013;7:1–3.

5. Scanlan PD, Stensvold CR, Rajilić-Stojanović M, Heilig HGHJGGHJG, De Vos WM, O’Toole PW, et al. The microbial eukaryote Blastocystis is a prevalent and diverse member of the healthy human gut microbiota. FEMS Microbiol Ecol. 2014;90:326–30.

6. Morton ER, Lynch J, Froment A, Lafosse S, Heyer E, Przeworski M, et al. Variation in Rural African Gut Microbiota Is Strongly Correlated with Colonization by Entamoeba and Subsistence. PLoS Genet. 2015;11:1–28.

7. Chudnovskiy A, Mortha A, Kana V, Belkaid Y, Grigg ME, Merad M, et al. Host-Protozoan Interactions Protect from Mucosal Infections through Activation of the Inflammasome. Cell. Elsevier; 2016;167:444–456.e14.

8. Brune A. Symbiotic digestion of lignocellulose in termite guts. Nat Rev Microbiol. 2014;12:168–180.

9. Glynn PW. Coral reef bleaching: ecological perspectives. Coral Reefs. 1993;12:1–17.

10. Stensvold CR, van der Giezen M. Associations between Gut Microbiota and Common Luminal Intestinal Parasites. Trends Parasitol. Elsevier Ltd; 2018;34:369–77.

11. Keeling PJ, del Campo J. Marine Protists Are Not Just Big Bacteria. Curr Biol. Elsevier Ltd; 2017;27:R541–9.

12. Parfrey LW, Walters WA, Lauber CL, Clemente JC, Berg-Lyons D, Teiling C, et al. Communities of microbial eukaryotes in the mammalian gut within the context of environmental eukaryotic diversity. Front Microbiol. 2014;5:1–13.

13. Wilmes P, Wampach L, Heintz-Buschart A, Hogan A, Muller EEL, Narayanasamy S, et al. Colonization and Succession within the Human Gut Microbiome by Archaea, Bacteria, and Microeukaryotes during the First Year of Life. Front Microbiol. 2017;8:1–21.

14. Wilcox JJS, Hollocher H. Unprecedented Symbiont Eukaryote Diversity Is Governed by Internal Trophic Webs in a Wild Non-Human Primate. Protist. Elsevier GmbH.; 2018;169:307–20.

15. Vestheim H, Jarman SN. Blocking primers to enhance PCR amplification of rare sequences in mixed samples - A case study on prey DNA in Antarctic krill stomachs. Front Zool. 2008;5:12.

16. Belda E, Coulibaly B, Fofana A, Beavogui A, Traore S, Gohl D, et al. Preferential suppression of Anopheles gambiae host sequences allows detection of the mosquito eukaryotic microbiome. Sci Rep. Springer US; 2017;in review:1–13.

17. Boessenkool S, Epp LS, Haile J, Bellemain EP, Edwards ME, Coissac E, et al. Blocking human contaminant DNA during PCR allows amplification of rare mammal species from sedimentary ancient DNA. Mol Ecol. 2012;21:1806–15.

18. Tan S, Liu H. Unravel the hidden protistan diversity: application of blocking primers to suppress PCR amplification of metazoan DNA. Appl Microbiol Biotechnol. Applied Microbiology and Biotechnology; 2018;102:389–401.

19. Wilcox TM, Schwartz MK, McKelvey KS, Young MK, Lowe WH. A blocking primer increases specificity in environmental DNA detection of bull trout (Salvelinus confluentus). Conserv Genet Resour. 2014;6:283–4.

20. Hino A, Maruyama H, Kikuchi T. A novel method to assess the biodiversity of parasites using 18S rDNA Illumina sequencing; parasitome analysis method. Parasitol Int. Elsevier B.V.; 2016;9–12.

21. Waidele L, Korb J, Voolstra CR, Künzel S, Dedeine F, Staubach F. Differential Ecological Specificity of Protist and Bacterial Microbiomes across a Set of Termite Species. Front Microbiol. 2017;8:1–13.

22. Lukeš J, Stensvold CR, Jirků-Pomajbíková K, Wegener Parfrey L, Parfrey LW. Are Human Intestinal Eukaryotes Beneficial or Commensals? PLOS Pathog. 2015;11:e1005039.

23. Bower SM, Carnegie RB, Goh B, Jones SR, Lowe GJ, Mak MW. Preferential PCR amplification of parasitic protistan small subunit rDNA from metazoan tissues. J Eukaryot Microbiol. 2004;51:325–32.

24. Yilmaz P, Kottmann R, Field D, Knight R, Cole JR, Amaral-Zettler LA, et al. Minimum information about a marker gene sequence (MIMARKS) and minimum information about any (x) sequence (MIxS) specifications. Nat Biotechnol. 2011;29:415–20.

25. Massana R, Gobet A, Audic S, Bass D, Bittner L, Boutte C, et al. Marine protist diversity in European coastal waters and sediments as revealed by high-throughput sequencing. Environ Microbiol. 2015;17:4035–4049.

26. Pernice MC, Giner CR, Logares R, Perera-Bel J, Acinas SG, Duarte CM, et al. Large variability of bathypelagic microbial eukaryotic communities across the world’s oceans. ISME J. 2016;10: 945–958.

27. Stoeck T, Bass D, Nebel M, Christen R, Jones MDM, Breiner H-W, et al. Multiple marker parallel tag environmental DNA sequencing reveals a highly complex eukaryotic community in marine anoxic water. Mol Ecol. 2010;19:21–31.

28. Amaral-Zettler LA, McCliment EA, Ducklow HW, Huse SM. A Method for Studying Protistan Diversity Using Massively Parallel Sequencing of V9 Hypervariable Regions of Small-Subunit Ribosomal RNA Genes. Langsley G, editor. PLoS One. 2009;4:e6372.

29. Gilbert JA, Jansson JK, Knight R. The Earth Microbiome project: Successes and aspirations. BMC Biol. 2014;12:1–4.

30. de Vargas C, Audic S, Henry N, Decelle J, Mahe F, Logares R, et al. Eukaryotic plankton diversity in the sunlit ocean. Science. 2015;348:1261605.

31. Comeau AM, Li WKW, Tremblay J-É, Carmack EC, Lovejoy C. Arctic Ocean microbial community structure before and after the 2007 record sea ice minimum. PLoS One. 2011;6:e27492.

32. Pawlowski J, Christen R, Lecroq B, Bachar D, Shahbazkia HR, Amaral-Zettler LA, et al. Eukaryotic Richness in the Abyss: Insights from Pyrotag Sequencing. Gilbert J, editor. PLoS One. 2011;6:e18169.

33. Klindworth A, Pruesse E, Schweer T, Peplies J, Quast C, Horn M, et al. Evaluation of general 16S ribosomal RNA gene PCR primers for classical and next-generation sequencing-based diversity studies. Nucleic Acids Res. 2013;41:1–11.

34. Cornejo-Tapia A, Orellana-Peralta F, Weilg P, Bazan-Mayra J, Cornejo-Pacherres H, Ulloa-Urizar G, et al. Etiology, epidemiology and clinical characteristics of acute diarrhea in hospitalized children in rural Peru. J Infect Dev Ctries. 2017;11:826.

35. Rognes T, Flouri T, Nichols B, Quince C, Mahé F. VSEARCH: a versatile open source tool for metagenomics. PeerJ. 2016;4:e2584.

36. Caporaso JG, Kuczynski J, Stombaugh J, Bittinger K, Bushman FD, Costello EK, et al. QIIME allows analysis of high-throughput community sequencing data. Nat Methods. Nature Publishing Group; 2010;7:335–336.

